# Estimation of complex effect-size distributions using summary-level statistics from genome-wide association studies across 32 complex traits and implications for the future

**DOI:** 10.1101/175406

**Authors:** Yan Zhang, Guanghao Qi, Ju-Hyun Park, Nilanjan Chatterjee

## Abstract

Summary-level statistics from genome-wide association studies are now widely used to estimate heritability and co-heritability of traits using the popular linkage-disequilibrium-score (LD-score) regression method. We develop a likelihood-based approach for analyzing summary-level statistics and external LD information to estimate common variants effect-size distributions, characterized by proportion of underlying susceptibility SNPs and a flexible normal-mixture model for their effects. Analysis of summary-level results across 32 GWAS reveals that while all traits are highly polygenic, there is wide diversity in the degrees of polygenicity. The effect-size distributions for susceptibility SNPs could be adequately modeled by a single normal distribution for traits related to mental health and ability and by a mixture of two normal distributions for all other traits. Among quantitative traits, we predict the sample sizes needed to identify SNPs which explain 80% of GWAS heritability to be between 300K-500K for some of the early growth traits, between 1-2 million for some anthropometric and cholesterol traits and multiple millions for body mass index and some others. The corresponding predictions for disease traits are between 200K-400K for inflammatory bowel diseases, close to one million for a variety of adult onset chronic diseases and between 1-2 million for psychiatric diseases.

## Introduction

Sample sizes for genome-wide association studies for many complex diseases and traits now range between tens to hundreds of thousands due to success of the large consortia. These studies have led to discoveries of dozens and sometimes hundreds of common susceptibility SNPs for individual traits^1–3^. Although the effects of individual markers are modest, collectively they provide significant insights into underlying pathways and contribute to models for risk-stratification for some common diseases such as breast cancer^4,5^. Existing GWAS for almost all traits also indicate that common variants have the potential to explain much more heritability than that explained by SNPs achieving stringent genome-wide significance level^6–13^.

It can be anticipated that sample sizes for many easily ascertainable traits and common diseases will continue to rise rapidly allowing GWAS to reach their full potential. However, for rare diseases and difficult or expensive-to-ascertain traits, it is not clear what is realistically achievable given the practical limits in sample size and what is the best way to distribute resources as a community based on the likely yield for the traits. We and others have earlier shown that yield of future GWAS critically depend on underlying effect-size distribution^14–16^. In this report, we propose novel methods for analysis of summary-level association statistics to estimate effect-size distributions and subsequently apply these methods across a large number of traits and diseases to make projections regarding future discoveries and genetic risk prediction.

Recently, LD-score method has become a popular approach for estimation of heritability and co-heritability using summary-level association statistics across GWAS^17–19^. The method relies on the observation that for highly polygenic traits, the association test-statistics for GWAS markers are expected to be linearly related to their *LD-scores*, a measure of total amount of LD individual SNPs have with others in the genome. The slope of this linear relationship is determined by the degree of narrow-sense heritability of the underlying trait associated with a reference panel of SNPs tagged by the GWAS markers. In this report, we develop a likelihood-based framework that allows estimation of potentially complex effect-size distribution of a trait based on a single set of summary-statistics that are widely available from GWAS consortia. We show that under a mixture-normal model for the effects of the genetic variants, the distribution of summary-statistics can be approximated by an alternative mixture normal distribution, the form of which depends on estimates of LD-scores and number of underlying tagged SNPs. We propose methods for obtaining estimates of model parameters and valid standard errors through a composite-likelihood inferential framework effectively dealing with correlated GWAS markers.

We apply the methods to analyze publicly available summary-level association statistics for 19 quantitative traits and 13 binary traits to provide the most comprehensive analysis of effect-size distributions underlying GWAS to date. The applications provide detailed insights into the diversity of genetic architecture of complex traits, including numbers of underlying susceptibility SNPs and different clusters of effects that contribute to heritability of the traits. Using these estimated effect size distributions, we then provide projections regarding yield of future GWAS, in terms of identification of susceptibility SNPs and in terms of building models for genetic risk prediction.

## Methods

### Data and Model

We assume data are available on estimated regression coefficients 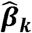 and corresponding standard error (**s**_**k**_) for ***k*= 1,…, *K*** GWAS markers assumed to be analyzed standardized scale so that genotypes and phenotypes have unit variances. Typically, these “summary-level” results are obtained from one-SNP-at-a-time “marginal” analyses that do not account for correlation across SNPs. As linear regression models can yield useful approximations for logistic and other non-linear regressions, it is useful to first describe the methods for the study of a continuous trait ***Y***. We assume that the GWAS markers tag a set of ***M*** SNPs in an underlying reference panel with respect to which a causal model can be defined in the form 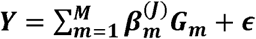. Here, the 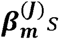 represent the regression coefficients in a “joint model” that accounts for correlation of the SNPs. The simple relationship 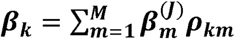, between regression coefficients of SNPs from a marginal model and those from a joint model, where the ***ρ*_*km*_***s* denote correlations across SNP-pairs, allows fitting of “joint” models from estimates of marginal regression coefficients^20^. The same relationship between joint and marginal effects approximately holds true for logistic regression models for relatively uncommon diseases.

We assume that the regression coefficients in the “joint model” are independently and identically distributed (*i.i.d.*) according to a mixture distribution in the form

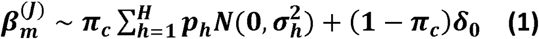

where, a fraction, 1 – π _c_ of the SNPs have no association with the trait. The model assumes the effect size distribution for non-null SNPs to be symmetric and modal around zero and allows distinct clusters of effect-sizes through incorporation of different variance component parameters 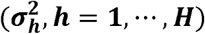. In our application, we consider fitting two-component (***M2***) or three-component (***M3***) models, which allow the distribution of association parameters for non-null SNPs to follow either a single normal distribution (***H*= 1**) or a mixture of two normal distributions (***H* = 2**). The latter model allows two distinct variance component parameters thereby allowing a fraction of SNPs (***p*_1_**) to have distinctly larger effects.

### Composite likelihood estimation

In principle, a joint likelihood for summary-level association statistics across GWAS markers can be derived using the relationship 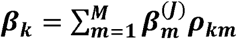 and the fact that 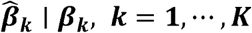 are expected to follow a multivariate normal distribution^21^. In general, however, the computation of this likelihood for genome-wide analysis of millions of SNPs can be complex. We show that under an assumption of independence of LD patterns and the probability of SNPs belonging to different mixture components in (1), the distribution of marginal effects for individual SNPs can be approximated by another mixture form (see **Supplementary Notes**)

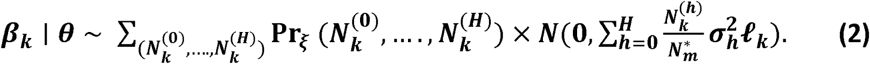

Here, 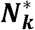 is the total number of SNPs in the reference panel in a “neighborhood” (***𝒩*_*k*_**) that may be “tagged” by marker *k*; 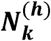 are latent variables indicating number of SNPs in ***𝒩*_*k*_** that have underlying effects from ***h*= 1,…, *H*** different components of the mixture distribution (see **Equation 1**); 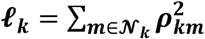 is the LD score for the *k*-th GWAS marker associated with 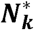 SNPs in the reference panel; ***ξ***= (***π***_**c**_, ***p*_1_**,…, ***p*_*H*-1_**) and 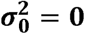. In the above formula, the mixing probability 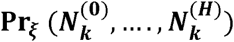 can be calculated based on the standard multinomial distribution with total counts defined by 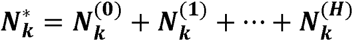 and cell probabilities given by (**1** - ***π*_*c*_**, ***π*_*c*_** × ***p*_*1*_**,…, ***π*_*c*_** × ***p*_*H*_**). Intuitively, (2) implies that the distribution of marginal effects of the GWAS markers is given by mixtures of mean zero normal distributions with variance component parameters determined by the product of LD-score and weighted sum of the variance component parameters of the original mixture model (see Equation 1). The weights which are defined by the number of different types of underlying effects a GWAS marker tags, are expected to follow a multinomial distribution in general and a binomial distribution in special cases where ***h***= 1.

If ***θ*** denotes the unknown parameter of model (1), we can write down the likelihood for an individual GWAS marker in the form

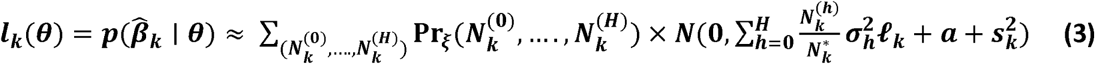

by exploiting the fact that 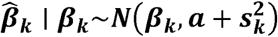, which incorporates an additional factor ***a*** that could account for systematic bias due to effects such as population-stratification. In computing (3), we exploit the fact that the number of underlying susceptibility SNPs 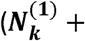 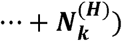 that may be tagged by an individual GWAS marker is likely to be small, e.g. less than or equal to 10, and so the number of terms in the mixture can be dramatically truncated to increase the speed of computation. To combine information across all markers, we propose forming a composite likelihood in the form 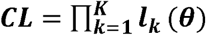, which ignores correlation in 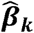 across SNPs. Following the theory of generalized estimating equations^22^, it is evident that such a composite likelihood approach will produce unbiased estimate of ***θ*** as long as (3) is a valid likelihood for the summary-statistics for the individual markers. We maximize the likelihood using an Expectation-Maximization algorithm, where in each M-step the mixing proportions (***π*_*c*_**, ***p*_1,_**… ***p*_*H*-1_**) are estimated in closed form and the variance component parameters are estimated by numerical optimization of weighted univariate normal-likelihoods **(see Supplementary Notes)**.

### Variance calculations

We obtain estimates of standard errors for parameter estimates based on a sandwich variance estimator associated with the composite likelihood (see **Equation 3**). Let ***s*_*k*_** (***θ***) = *∂logI*_**k**_(***θ***)/*∂****θ*** denote the score function associated with the likelihood ***I*_*k*_**(***θ***) for the *k*th GWAS marker and let 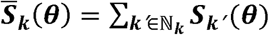 be the total likelihood-score across all the GWAS markers that are in the neighborhood of the *k*th marker, including itself, i.e. 𝒩_***k***_. It is important to note that unlike the calculation of the total LD-score that involves SNPs in the underlying reference panel, the total likelihood score is computed only with respect to the set of markers that are included in the GWAS study itself. Further, we define 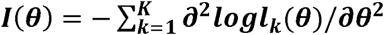 to be the total information matrix associated with the composite likelihood. The sandwich variance estimator^23,24^ is now defined as

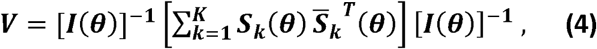

which itself can be estimated by plugging in the estimated parameter values 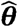 in lieu of ***θ***. The estimator accounts for correlation across the GWAS markers through calculation of empirical variance-covariances across the likelihood-scores within sets of correlated markers defined by physical distance and LD-thresholding criterion same as the one used to define the LD-scores. The estimator is expected to produce valid estimates of standard errors for the parameter estimates even when the underlying model is misspecified.

### Calculation of LD-score (l_k_) and number of tagged SNPs 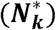

To implement the proposed method, we need to estimate the number of underlying SNPs in the reference panel tagged by the GWAS markers and the corresponding LD-scores. As we analyze GWAS of primarily Caucasian studies, we obtain required information based on analysis of 489 individuals of European origin from the 1000 GENOME project Phase 3 study^25^. We evaluated all the LD-scores and number of tagged SNPs based on a reference panel of ∼1.2 million common SNPs that were included in the Hapmap3 panel. We estimated 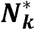 by the number of SNPs in the reference panel which are within 1Mb distance and have an estimated LD coefficient with the GWAS marker above a fixed threshold (e.g. ***r*^*2*^**> 0.1). Then we calculate the corresponding LD-score by summing up the corresponding squared LD coefficients. We evaluate sensitivity of our results with respect to variation in ***r*^*2*^** thresholds. In calculation of LD-score, we employ the same bias-correction adjustment which is used in the LD-score regression^17^.

Across all traits, we first extract association statistics available from the underlying GWAS for the set of the Hapmap 3 SNPs (see **Supplementary Table 1** for list of data sources). As most studies provide results after imputation, association statistics are available for large majority of the Hapmap 3 SNPs across these studies. We then follow the same filtering steps as in LD-Hub^19^ to select GWAS markers to standardize the summary-level datasets. In particular, we include SNPs with MAF above 5% and remove SNPs that had sample sizes less than 0.67 times the 90^th^ percentile of sample sizes or were within the major histocompatibility complex (MHC) region (i.e., SNPs between 26Mb and 34Mb on chromosome six), or had extremely large effects (*Χ* **^2^**> 80).

### Simulation Studies

We use a novel simulation scheme to generate summary-level association statistics for GWAS without generating individual-level data under a given model for genetic architecture of the trait. We simulate summary-statistics based on the model

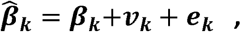

where ***v*_*k*_**s are assumed to be *i.i.d.* following normal distribution with mean zero and variance *a* and 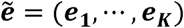 is assumed to be following a multivariate normal distribution with mean zero and variance-covariance matrix ***R/n***, with ***n*** denoting the sample size for GWAS and 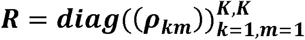 being the matrix of LD coefficients across the GWAS markers. In each simulation, we first generate value for 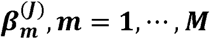, ***m = 1,…, M*** based on model (1) and then generate values for, ***β***_*k*_***,K*** = **1,…, *K*** based on the transformation 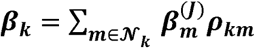. For simulation of 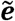, we observe that in a GWAS study where the phenotype has no association with any of the markers, the summary-level association statistics is expected to follow the same multivariate distribution as 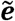. Thus, we simulate null phenotypes for the samples in our reference dataset and calculated association statistics 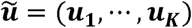 for the GWAS markers. We then define 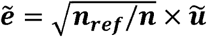 to account for the difference in sample sizes between the reference dataset and the GWAS.

## Results

Simulation studies show that the proposed method produces nearly unbiased estimates of parameters and associated standard errors when data are both simulated and analyzed using the two-component model (**Supplementary Table 2**). However, when data are simulated using the three-component model (**M3**), but analyzed using the simpler two-component model (***M2***), the estimates of proportion of susceptibility SNPs (***π*** _**c**_) and variance component parameter (**σ ^2^**) are downwardly and upwardly biased, respectively. The estimates of total heritability and estimates of standard errors for all parameters are nevertheless nearly unbiased. When simulation and analysis are, both conducted using the three-component model, the degree of bias in estimates of ***π*** _**c**_ reduces to a large extent and continues to decrease with increasing sample size. Across all sample sizes, the three-component model estimates distinct variance component parameters associated with the two components for the non-null effects. For studies with smaller sample sizes (i.e. *N* < 25*K*), there is substantial bias and uncertainty in estimates of cluster specific variance component parameters and mixing proportions. Despite such uncertainty, uniformly across all sample sizes, the three-component model provides more accurate estimate of number of susceptibility SNPs in the tails regions of effect-size distribution (**Supplemental Table 3**).

We analyze summary-level results from GWAS for each trait using both the two- and three-component models and assess their goodness of fit of by comparing observed distribution of p-values against what is expected based on fitted models (see **Online Methods**). In general, the three-component model provides distinctly better fit for the observed distribution of p-values (**Figure 1, Supplementary Figures 1-5**). For most of the traits, it provides excellent to adequate fit to observed p-values over a wide range except at extreme tails of the distribution (p-value < 10^-10^) indicating the presence of a small number of susceptibility SNPs whose underlying effects were “outliers” with respect to the fitted effect-size distribution. Interestingly, for a subset of the traits, consisting of psychiatric diseases and traits related to intelligence, educational accomplishments and cognitive ability, the fits for the two- and three-component models are very similar indicating that the effect-sizes for underlying susceptibility SNPs can be adequately modeled using a single normal distribution.

**Figure 1.**
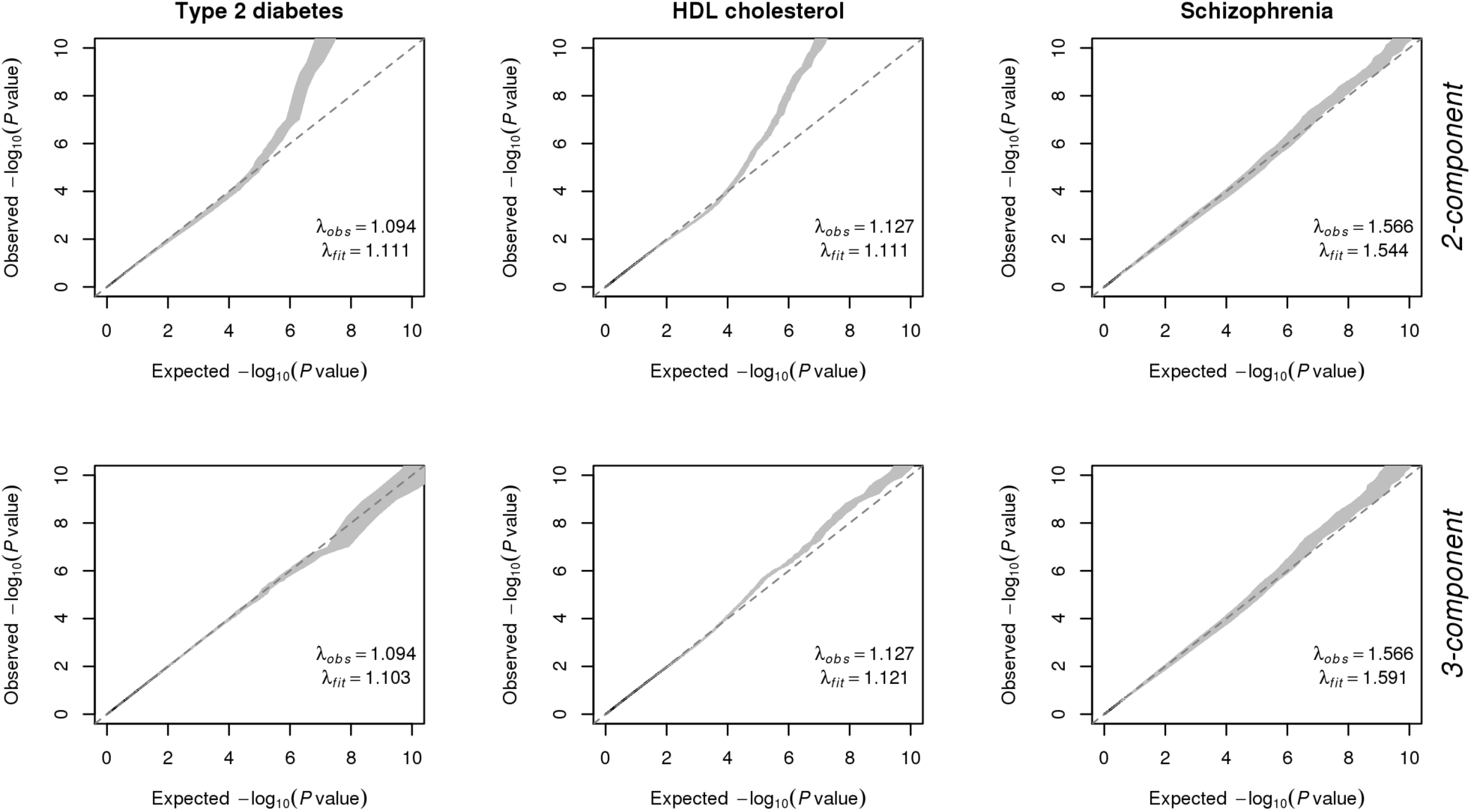
Q-Q plots comparing observed distributions of association statistics against those expected under fitted models for three representative traits. Plots in upper and lower panels are generated under 2- and 3-component models for underlying effect-size distributions, respectively. Shaded regions mark 80% point wise confidence intervals derived from 100 simulations (see **Methods** and **Supplemental Notes**). While the more flexible 3-component model provides distinctively better fit for Type 2 diabetes and HDL cholesterol, the simpler 2-component model is adequate for Schizophrenia. See **Supplementary Figures 1-5** for analogous plots for 29 additional complex traits.

Parameter estimates associated with the three-component model reveal wide diversity in genetic architecture across the traits (**Table 1, Figure 2, Supplementary Table 4**). Estimates of narrow sense heritability from the fitted models are generally close to those reported by LD-score regression. Estimates of the number of underlying susceptibility SNPs also vary widely, sometimes even among traits with similar estimates of heritability. In general, anthropometric traits, psychiatric diseases and traits related to education ability and cognitive performance are found to be most polygenic, each involving often close to 10,000 or even higher number of underlying susceptibility SNPs. In contrast, some of the early growth traits, autoimmune disorders and adult onset chronic common diseases (e.g., Coronary artery disease, Asthma, Alzheimer disease, Type-2 diabetes) are less polygenic, although each still involved at least a few thousands of underlying susceptibility SNPs. Consistent with results from simulation studies, we observe that the fitting of two-component model generally provides substantially lower estimate for the number of susceptibility SNPs (**Supplementary Table 5**).

**Figure 2.**
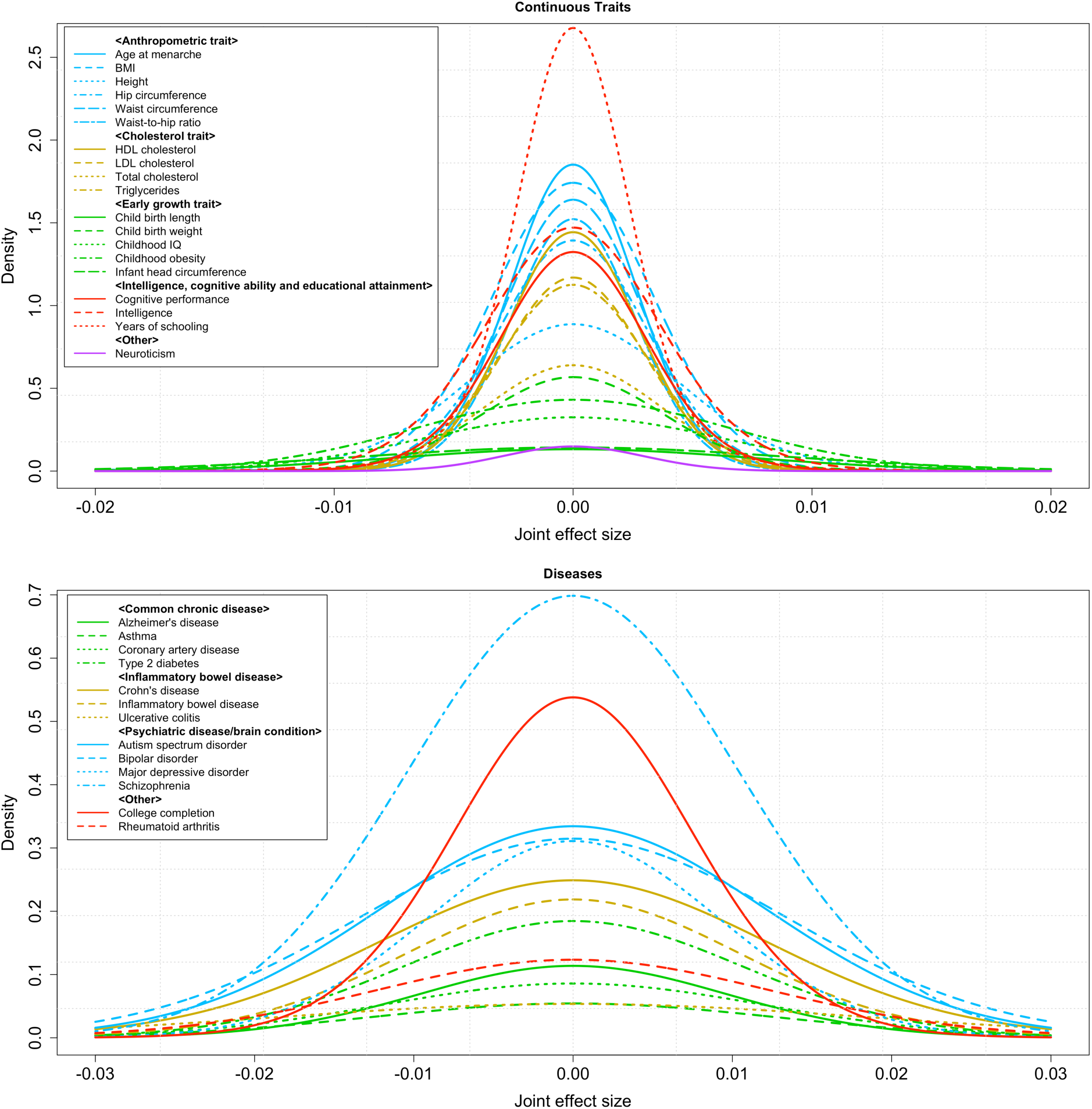
Estimated effect-size distributions based on fitted 3-component models to results from GWAS for continuous (upper panel) and binary traits (lower panel). The density plot for each trait is obtained based on mixture normal distribution where SNPs with null effects were represented by normal distribution with extremely small variance component 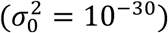. Distributions with fatter tails imply the underlying traits have relatively larger number of susceptibility SNPs with larger effects. In general, traits related to mental health and ability have effect-sizes with narrower trails in spite of larger estimate of heritability and associated number of susceptibility SNPs. See **Supplementary Table 6** for more detailed comparison of the tail regions of effect-size distributions.

**Table 1.** Estimated parameter values and standard errors from fitting of 3-component models for effect size distributions across 32 traits. All results are reported with respect to a reference panel of 1.2 million common SNPs included in the Hapmap3 panel after removal of MHC region. A *r*^”^threshold of 0.1 is used to define the set of reference SNPs the GWAS markers may tag. For traits related to mental health and ability, the variance component parameters associated with two non-null components of the mixture distribution collapsed indicating adequacy of a two-component model. Results from 2-component model and alternative *r*^”^threshold are shown in **Supplemental Tables 5 and 7.**

| Trait | Sample size (in 10K) | Total # of sSNPs<sup>a</sup> (SE<sup>b</sup>), reported in hundreds | # of sSNPs in cluster 1 (SE), reported in hundreds | Heritability explained by cluster 1<sup>c</sup> (SE) | Heritability explained by cluster 2<sup>d</sup> (SE) | Total heritability<sup>e</sup> (SE) |
| --- | --- | --- | --- | --- | --- | --- |
| **Continuous Traits:** |  |  |  |  |  |  |
| BMI | 12.4 | 174 (19) | 0.9 (0.34) | 0.02 (0.005) | 0.20 (0.012) | 0.21 (0.011) |
| Height | 13.4 | 120 (14) | 10.2 (1.65) | 0.15 (0.017) | 0.21 (0.018) | 0.36 (0.016) |
| Hip circumference | 21.3 | 130 (15) | 3.0 (1.07) | 0.02 (0.006) | 0.12 (0.010) | 0.15 (0.008) |
| Waist circumference | 23.2 | 140 (15) | 2.0 (0.72) | 0.02 (0.004) | 0.11 (0.008) | 0.13 (0.007) |
| Waist-to-hip ratio | 21.2 | 118 (17) | 2.5 (1.32) | 0.02 (0.005) | 0.08 (0.009) | 0.09 (0.007) |
| HDL cholesterol | 9.5 | 118 (15) | 2.1 (0.54) | 0.03 (0.005) | 0.09 (0.011) | 0.12 (0.011) |
| LDL cholesterol | 9.5 | 104 (19) | 1.4 (0.42) | 0.03 (0.005) | 0.09 (0.012) | 0.12 (0.011) |
| Total cholesterol | 9.5 | 71 (19) | 1.7 (0.47) | 0.04 (0.007) | 0.09 (0.011) | 0.13 (0.012) |
| Triglycerides | 9.5 | 104 (14) | 0.7 (0.20) | 0.02 (0.003) | 0.10 (0.011) | 0.12 (0.011) |
| Child birth length | 2.8 | 29 (12) | N/A<sup>f</sup> | N/A | N/A | 0.16 (0.034) |
| Child birth weight | 14.4 | 65 (24) | 3.3 (3.21) | 0.03 (0.018) | 0.09 (0.018) | 0.12 (0.009) |
| Childhood obesity | 1.4 | 84 (25) | 0.6 (0.24) | 0.05 (0.021) | 0.35 (0.058) | 0.41 (0.056) |
| Infant head circumference | 1.1 | 38 (23) | N/A | N/A | N/A | 0.30 (0.074) |
| Childhood IQ | 1.2 | 60 (33) | N/A | N/A | N/A | 0.24 (0.072) |
| Cognitive performance | 10.7 | 121 (43) | N/A | N/A | N/A | 0.12 (0.012) |
| Intelligence | 7.8 | 161 (81) | N/A | N/A | N/A | 0.24 (0.025) |
| Years of schooling | 29.4 | 194 (28) | 39.2 (32.57) | 0.07 (0.037) | 0.08 (0.037) | 0.15 (0.006) |
| Age at menarche | 18.2 | 146 (15) | 4.0 (0.59) | 0.05 (0.006) | 0.10 (0.007) | 0.15 (0.008) |
| Neuroticism | 16.1 | 13 (9) | N/A | N/A | N/A | 0.01 (0.005) |
| **Disease Traits:** |  |  |  |  |  |  |
| Alzheimer's disease | 1.7/3.7<sup>g</sup> | 33 (31) | 0.5 (0.48) | 0.09 (0.047) | 0.30 (0.080) | 0.39 (0.081) |
| Asthma | 1.0/1.6 | 20 (16) | 0.3 (0.11) | 0.15 (0.055) | 0.32 (0.120) | 0.47 (0.132) |
| Coronary artery disease | 2.2/6.5 | 30 (11) | 0.2 (0.11) | 0.03 (0.014) | 0.43 (0.072) | 0.46 (0.075) |
| Type 2 diabetes | 1.2/5.7 | 59 (33) | 0.5 (0.47) | 0.09 (0.046) | 0.67 (0.111) | 0.76 (0.111) |
| Crohn's disease | 0.6/1.5 | 92 (19) | 3.9 (0.83) | 1.49 (0.212) | 1.31 (0.267) | 2.80 (0.296) |
| Inflammatory bowel disease | 1.3/2.2 | 71 (29) | 4.2 (0.75) | 0.91 (0.122) | 0.72 (0.145) | 1.63 (0.159) |
| Ulcerative colitis | 0.7/2.0 | 30 (18) | 2.0 (0.91) | 0.60 (0.185) | 0.97 (0.224) | 1.57 (0.230) |
| College completion | 2.2/7.3 | 120 (69) | N/A | N/A | N/A | 0.73 (0.067) |
| Rheumatoid arthritis | 1.4/4.4 | 46 (18) | 1.8 (0.58) | 0.32 (0.063) | 0.68 (0.090) | 1.00 (0.098) |
| Autism spectrum disorder | 0.5/0.5 | 120 (88) | N/A | N/A | N/A | 1.77 (0.528) |
| Bipolar disorder | 0.7/0.9 | 124 (68) | N/A | N/A | N/A | 2.22 (0.272) |
| Major depressive disorder | 0.9/1.0 | 84 (40) | N/A | N/A | N/A | 0.71 (0.172) |
| Schizophrenia | 3.4/4.3 | 213 (27) | N/A | N/A | N/A | 2.28 (0.101) |

<sup>a</sup>Susceptibility SNPs. <sup>b</sup>Standard errors. <sup>c</sup>Cluster 1 and <sup>d</sup>cluster 2 refer to the two components for the normal-mixture model for effect-sizes for non-null SNPs where cluster 1 corresponds to the component with larger variance-component parameter. <sup>e</sup>Total heritability is defined as $h^2 = M\pi_c\{p_1\sigma_1^2 + (1 - p_1)\sigma_2^2\} = h_1^2 + h_2^2$ , where $M$ is the total number of SNPs in the Hapmap3 panel, $\pi_c$ is the proportion of susceptibility SNPs, $p_1$ is the proportion of SNPs in the 1<sup>st</sup> cluster among all the susceptibility SNPs, and $\sigma_1^2$ and $\sigma_2^2$ are the variance estimates corresponding to each cluster – this definition corresponds to heritability in observed scale for continuous traits and in log-odds-ratio scale for disease traits. <sup>f</sup>N/A implies the variance component parameters associated with two clusters collapsed for these traits and thus results are reported assuming existence of only one cluster of non-null SNPs. <sup>g</sup>Number of cases/ number of controls.

For a majority of the traits, the fitting of three-component model detects presence of distinct clusters of effects. For these traits, the average heritability explained per variant in one cluster 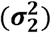 is often 10-fold or higher than that 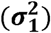 in the other cluster. Although a small fraction, typically ranging between 0.5-10%, of the susceptibility SNPs belong to clusters with larger effect-sizes, the fraction of heritability they explained is substantial for many traits (i.e. 10-50%). In contrast, for all psychiatric diseases and traits related to intelligence and cognitive ability, the estimates of two variance components collapse to a single value, a phenomenon consistent with adequate fit for the two-component model for these traits (see **Figure 1, Supplementary Figure 1-5**). Comparison of the number of SNPs in the tail regions of effect-size distribution show that some traits, such as early growth traits and inflammatory bowel diseases, have distinctly larger number of SNPs with moderate to large effects (i.e. odds-ratio > 1.03) (see **Supplementary Table 6**).

The diversity of genetic architecture across the traits implies major differences in the future yield of GWAS (**Figure 3**). In general, the expected number of discoveries will continue to rise rapidly across all traits in the foreseeable future. The degree of genetic variance they will explain will rise at a slower rate as the effect-size explained per SNP will continue to diminish. For most quantitative traits, the rate of increase in genetic variance explained is expected to diminish after sample size reaches approximately 300K. In contrast, for BMI, years of education and neuroticism, the genetic variance explained is expected to increase at a steady rate until sample size reaches at least one million. The sample sizes needed to identify SNPs that can explain 80% of GWAS heritability vary from 300K-500K for some of the early growth traits, between 1-2 million for some anthropometric and cholesterol traits and into multiple millions for the remaining. For most disease traits, genetic variance explained is expected to rise either steadily, or even at an accelerated rate for the highly polygenic psychiatric diseases, between sample sizes 50K-300K. The sample sizes needed to identify SNPs that can explain 80% of GWAS heritability turn out to be between 200K-400K for inflammatory bowel diseases, around 600K for rheumatoid arthritis, around one million for most common adult onset chronic diseases and between 1-2 million for psychiatric diseases.

**Figure 3.**
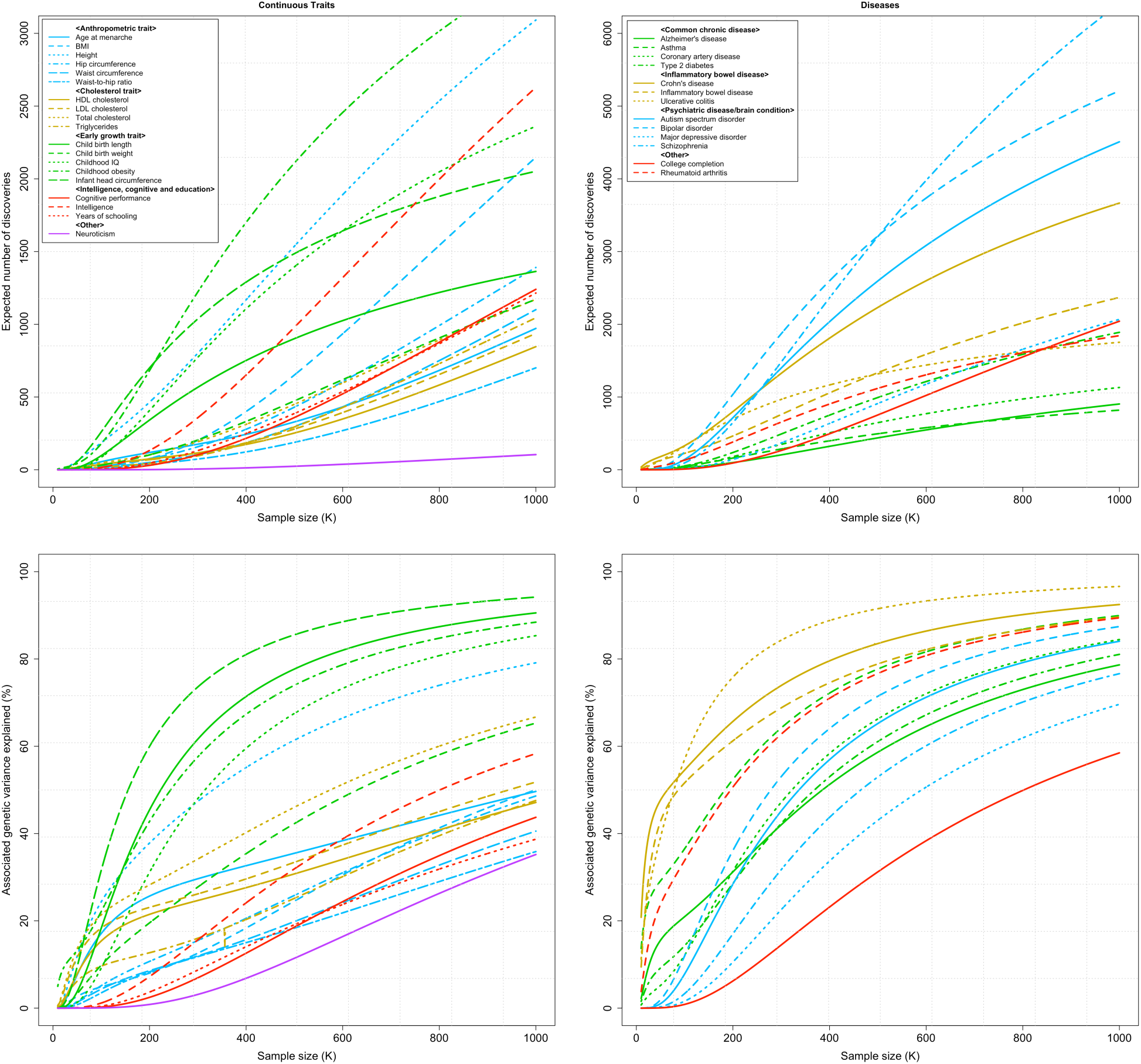
Projected number of discoveries (upper panel) and corresponding percentage of GWAS heritability explained (lower panel) based on fitted 3-component models for effect-size distribution for continuous (left panel) and binary traits (right panel). Results are based on power calculations for discovery at the genome-wide significance level (p-value=5×10^−8^). Total estimate of GWAS heritability is used as the denominator in calculating the percentage of heritability that would be explained by SNPs reaching genome-wide significance.

We conduct several sensitivity analyses to ascertain the robustness of results. We observe that use of higher ***r*^2^** threshold for defining tagging SNPs lead to increase in estimate of number of susceptibility SNPs across all traits though conclusions regarding relative degrees of polygenicity of the traits and existence or absence of distinct clusters of effects do not alter (**Supplementary Table 7**). To determine impact of sample size, we used three traits namely, *years of education, child-birth-weight and schizophrenia*, for which we accessed summary-level statistics available from older GWAS with sample sizes markedly smaller (by 3-5 fold) than the more recent studies we have analyzed. For all the three traits, the estimated number of susceptibility SNPs increased in the more recent studies, with the increase being particularly prominent for years of education and child-birth-weight (**Supplementary Table 8**). These results are consistent with simulation studies where we observe that the estimates tend to increase towards true value as sample size increases (see **Supplementary Table 2**). Thus, it is likely that the true genetic architecture of these traits is likely to be even more polygenic than that suggested by our analysis.

We also attempt to empirically validate the ability of the fitted effect-size models to project yields of future studies (**Supplementary Table 9**). In this analysis, we include two additional traits namely, *height* and *BMI*, for both of which results are available from more recent studies than the ones we used to build the models. For all traits, except *years of education*, the fitted model built based on older studies are able to predict the genomic control factor for the newer study reasonable well. For three of the traits namely, *Height, BMI* and *schizophrenia*, the model also predict accurately the number of independent loci reaching genome-wide significance level. For *years-of-education* and *child-birth-weight*, the model over-predicts, the expected number of independent discoveries.

Using the inferred effect-size distributions and theoretical framework we developed earlier^16^, we assess the expected predictive performance of polygenic models when SNPs are included at optimal threshold^26^ and the stringent genome-wide significance level (p-value <5 x 10^-8^). The results reveal two very distinct patterns. For psychiatric diseases, which include a continuum of highly polygenic effects, use of the optimal threshold for SNP selection is expected to lead to large improvement in performance of polygenic models in a wide range of sample sizes (**Figure 4, and Supplementary Figure 6**). For these traits, the optimal threshold is expected to be highly liberal (i.e. p-value > 0.2) for relatively small studies (i.e. **n** < 20,000) and then become more stringent as sample size increases. In contrast, for all other diseases, which are less polygenic but included more SNPs with relatively large effects, use of optimal threshold is expected to lead to only modest benefits. For these diseases, the optimal threshold is expected to be highly stringent for studies with small sample sizes (**n** < 10*K*), then gradually become more liberal with intermediate sample sizes (10K < **n** < 50*K*) and then slowly decrease for larger sample sizes.

**Figure 4.**
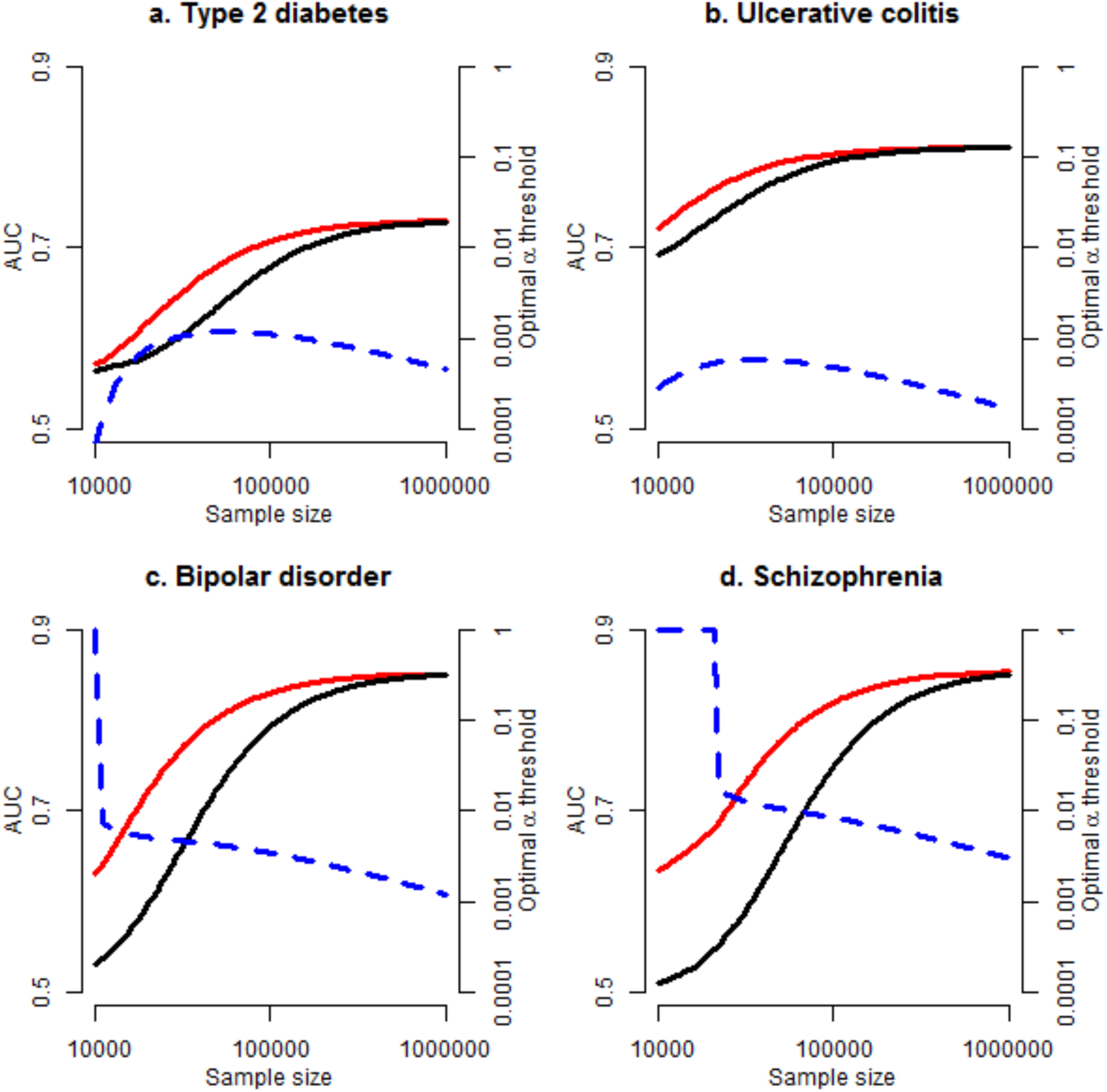
Expected area under the curve (AUC) for polygenic risk prediction models with SNPs included at the optimal significance (α) threshold (red solid line) and at the genome-wide significance level of 5×10^−8^(black solid line). Optimum values for significance thresholds (shown in blue dashed line) are obtained based on expected relationship of AUC with sample size and significance threshold under the fitted 3-component models for effect-size distributions. All calculations assume an analysis of a total of 200,000 independent set of SNPs.

We also assess the potential implications for the inferred effect-size distributions on subsequent analysis of GWAS to optimize SNP discovery and estimation of SNP effect-sizes. We calculate local false discovery rates^27,28^ for each SNP based on the observed Z-statistics and the model for the marginal effect-sizes (see **Equation 2**). It is evident that a high degree of polygenicity for the traits implies that it may be possible to identify large number of loci at fairly low false discovery rates (**Supplementary Figures 7-8**). The posterior mean estimates for effect-sizes for individual SNPs are shrunken heavily towards zero compared to their estimates available from GWAS with the degree of shrinkage being highest for SNPs with intermediate effects and studies with smallest sample sizes (**Supplemental Figure 9-10**). Further, SNPs with largest effect-sizes are shrunken more under the two-component than the three-component model because of the ability of the latter model to accommodate SNPs with distinctly larger effects.

## Discussion

In this report, we develop and apply novel methods for analysis of GWAS summary-level statistics to conduct most comprehensive analysis of effect-size distributions associated with common genetic variations across complex diseases and traits. Our analysis shows that fairly parsimonious normal-mixture models for effect-size distributions, involving four or less parameters, can adequately describe distribution of p-values observed in some of the most recent and largest GWAS. The estimated values of the model parameters provide insights into heritability, degree of polygenicity and presence of distinct clusters of effects. Using these estimated effect-size distributions, we provide further insights into likely yields of common variant GWAS in future, as sample size further accumulates.

Estimation of heritability based on SNP-arrays has been a major focus of research for GWAS ever since the first application of the approach to analysis of human height^29–33^. While such estimates of heritability provide an understanding of the limits of GWAS to explain trait variations, further understanding of effect-size distribution is critical for understanding how fast one can approach the limit as a function of the corresponding sample-size^15,16^. Although applied in more limited settings, various approaches have been developed in the past for estimation of effect-size distribution (ESD) from GWAS. We have described a simple method for estimating ESD within the range of effects observed in an existing study simply based on its reported number of findings with different effect sizes and its power for discovery of the underlying study at those effect-sizes^14^. Several methods have been developed to infer ESD by evaluating predictive performance of series of polygenic models that include varying number of SNPs on independent validation datasets^34,35^. A variety of Bayesian methods described for analysis of GWAS studies can also produce estimates of effect-size distribution according to the underlying “prior” models^36–40^. Most recently, a number of methods have been proposed for analyzing GWAS summary-level data under the two-component mixture model for estimating effect-size distribution^21–41.42^.

The current analysis of effect-size distribution is unique in several ways. First, we provide most comprehensive insights into diversity in effect-size distributions across complex traits by analysis of publicly available summary-level statistics from a large number of GWAS. Second, by considering a flexible model for effect-size distribution, we show that a commonly used two-component model, which assumes that the effect-sizes for underlying causal SNPs can be described by a single normal distribution centered around zero, can be inadequate for describing effect-size distribution across a large majority of the traits. Instead, a three-component model for effect-size distribution, which allows a proportion of causal SNPs to have distinctively larger effects than others, provides adequate fit to current GWAS for most traits and is thus likely to provide more accurate projections for future discoveries. Further, through both simulation studies and data analysis we demonstrate that there could be additional hidden components of effect-size distributions associated with groups of causal SNPs with extremely small effects that have no power to be distinguished from null effects in current studies.

In terms of methodology, the proposed approach, although is closely related to several recent methods^21,42^, has some unique aspects. We show that under the commonly invoked assumption of independence of effect-sizes and local linkage disequilibrium pattern, the likelihood of summary-statistics for individual GWAS markers depends on LD coefficients through total LD-score. The simplification allowed us to develop a computationally tractable and robust method for estimating parameters, as well as their standard errors, under the complex mixture model for effect-size distribution based on an underlying composite likelihood inferential framework. Simulation studies show that the proposed method produces unbiased estimate of heritability even when the underlying model of effect-size distribution is misspecified. Further, it provides either nearly unbiased or downwardly biased estimate of proportion of non-null SNPs depending on sample size and whether the number of components for mixture model is correctly or not. The simulation study also shows that the proposed standard error estimator is highly accurate and thus could be used to conduct statistical inference.

Estimation of the total number of susceptibility SNPs for an underlying trait can be an elusive concept given the sensitivity of the quantity to underlying model assumptions and sample size of GWAS. Given that any analysis of effect-size distribution can only be expected to provide a lower bound of the total number of susceptibility SNPs, the possibility of infinitesimal^43,44^ or omnigenic model^45^ where every SNP or gene is associated with a trait cannot be ruled out. Given such difficulty, a more interpretable way to compare genetic architecture and discovery potential of GWAS across traits would be to examine the number of susceptibility SNPs that may have meaningfully large effects, such as an odds-ratio of 1.01 or larger for disease traits (**Supplemental Table 6**).

Our projections show that high-degree of polygenicity of the traits imply requirement of very large sample sizes, from hundreds of thousands to millions, for discovery of SNPs at genome-wide significance level in order to explain nearly all of the GWAS heritability. These projections are expected to be optimistic given that larger studies in the future with increasingly heterogeneous sample may reveal higher degree of polygenic nature of the underlying traits. However, given that thousands to tens of thousands of SNPs may be associated with any individual trait, the current practice of using stringent genome-wide significance level to minimize the chance for a single false positive may be too conservative an approach for discovery. Instead, more optimal strategy for discovery would be to select thresholds in a more adaptive fashion, taking into account underlying effect-size distributions, while controlling for false discovery rate^46,47^.

A major utility of future GWAS could be improving performance of polygenic prediction model as opposed to simply identifying of susceptibility SNPs at high-levels of significance^48–52^. Our projections show that use of optimal thresholds will lead to large benefit for psychiatric diseases, but much more moderately so for others. In general, across all traits we observe that the overall discriminatory performance of models, as measured by the area under the curve (AUC) criterion, is expected to rise very modestly after sample size reaches around 100K. However, larger sample size could still improve the performance of models meaningfully in terms of identifying individuals who are at extremes of risk distribution. For example, for type-2 diabetes, a model built on GWAS with sample size of 1 million instead of 100K individuals is expected to identify additional 0.2% (1.2% vs 1.4%) of the population who are at 5-fold or higher risk than the average risk of the general population. Such improved model may lead to intervention for an additional 2.2% of prospective cases (8.2 vs 10.4%) (**Supplementary Table 10**).

The limitations of the proposed method include that its assumption of independence of effect-sizes from allele frequencies and local LD patterns of the SNPs. It has been recently shown that these simplified assumptions, which implicitly or explicitly have been used in many earlier methods, can lead to substantial underestimation of heritability^53^ and is likely to have impact on other aspects of effect-size distributions as well. In principle, the proposed method can be extended to model dependence of effect-sizes on various SNP characteristics through regression modeling approach. The proposed inferential framework, which yields parameter estimates as well as their standard errors, can be used to test various hypotheses regarding underlying parameters of such models and thus could provide insights into genetic architecture of traits in finer details. It will also be of interest to extend the framework to jointly model the effect-size distributions to obtain deeper insights into shared genetic architecture of multiple traits.

To summarize, we propose methods for statistical inference for effect-size distributions under flexible normal-mixture models using summary-level GWAS statistics. Applications of the methods to a large number of GWAS reveal wide diversity in genetic architecture of the underlying traits with important consequence for the yields of future GWAS in terms of both discovery and risk prediction.

## Authors Contribution

Y.Z. and N.C. conceived the methods. Y.Z., G.Q. and J.P. carried out all analyses. Y.Z. and N.C. wrote the manuscript. All authors reviewed the manuscripts.

## Acknowledgement

The authors would like to thank Haoyu Zhang for assistance with computing and Manjushree Chatterjee for editing of manuscript. The research was supported by Bloomberg Distinguished Professorship endowment.

